# Longitudinal Plasma Proteome Changes Before and After Catheter Ablation in Atrial Fibrillation

**DOI:** 10.64898/2026.01.07.698107

**Authors:** Chi Him Kendrick Yiu, Chenhan Sam Ma, Tim R. Betts, Kim Rajappan, Matthew Ginks, Michala Pedersen, Yaver Bashir, Rohan Wijesurendra, Svetlana Reilly

## Abstract

**Background:** Proteins in human plasma serve as critical markers for predicting disease risk and guiding therapeutic development. Current prediction models for atrial fibrillation (AF) largely rely on electronic health records; however, the plasma proteome of patients with AF reflects key biological processes, including inflammation, that are not captured by clinical variables alone. Circulating inflammatory mediators contribute to electrical and structural remodelling in the atria, thereby sustaining the AF phenotype. Identification of plasma proteins associated with AF may therefore improve understanding of the inflammatory and other biological processes underlying AF pathophysiology.

**Objective:** In this study, we profiled the plasma proteome of patients with paroxysmal AF (pxAF), persistent AF (persAF), and non-AF controls using Olink assay technology.

**Methods:** Plasma samples from 30 individuals were analysed with the Olink Reveal panel. Differential expression analysis of normalised protein expression (NPX) values was performed between groups, with differentially expressed proteins (DEPs) defined by *P* < 0.05.

**Results:** We identified 87 DEPs in pxAF and 107 DEPs in persAF compared with controls. From these, we shortlisted 11 candidate proteins that were upregulated in persAF at baseline and showed reduced expression 12 months after catheter ablation. This subset of proteins is implicated in the regulation of inflammation (CCL23, CXCL10, IL33), metabolism (ALDH3A1, NDUFS6), cell-matrix adhesion (AFAP1L1, LGALS7, SPOCK1), and physiological signalling (NOS1, PROK1, PTH).

**Conclusion:** Collectively, these plasma proteins highlight systemic molecular mechanisms contributing to AF pathogenesis and represent potential AF-specific biomarkers warranting further investigation in larger clinical cohorts and mechanistic studies.

## Introduction

Circulating proteins in the human body play important roles in reflecting physiological and pathophysiological processes. In particular, the plasma proteome offers opportunities to explore new biomarkers and therapeutic targets, improve disease risk prediction, and provide better mechanistic insights ^1^. Measuring proteins in human plasma can be promising as a minimally invasive diagnostic approach in precision medicine ^2^. Additionally, proteins serve as better candidates than genes as they reflect the ultimate biological activity and functions, while the complex transcriptional regulatory processes of genes increase the uncertainties in mechanistic understanding of gene-disease relationships. However, limitations persist in studying clinically relevant protein markers due to the lack of high-throughput and sensitive assays to measure protein abundance, in which accurate disease prediction is typically challenging for lowly expressed proteins ^3^.

Atrial fibrillation (AF) remains a major public health concern with a global prevalence of 52.55 million individuals ^4^. As the most prevalent type of arrhythmia, AF is closely associated with an elevated risk of cardiovascular diseases (e.g., stroke, myocardial infarction, heart failure), cognitive impairment, and dementia ^5^. When patients have AF coexisting with these complications, it results in increased mortality rate ^6^. Development of AF prediction models using machine learning is an emerging field ^7–11^, but most of these models were established based on physiological and clinical features extracted from electronic health records. Moreover, prediction of incident AF has been the primary focus of these models, while the importance of other classifications such as paroxysmal and long-lasting AF has often been overlooked. Accurate prediction of these AF phenotypes may be hindered by underlying heterogenous biological mechanisms, resulting in complex electrical and structural remodelling ^12^. Hence, there is an obvious neglect of the significance of using circulating protein markers in AF prediction models and integrating them with existing electronic health records may be beneficial to improve our pathophysiological understanding of AF. In addition to conventional biomarkers of AF such as natriuretic peptides, cardiac troponins, and C-reactive protein (CRP), other studies have identified emerging markers of inflammation and myocardial damage that are elevated and associated with worsened cardiovascular outcomes in AF patients, including interleukin-6, growth differentiation factor-15 and insulin-like growth factor-binding protein-7 ^13,14^. Proteomic studies have also highlighted extracellular matrix organisation and inflammation-related pathways among proteins associated with incident AF^15^.

Inflammation is a crucial biological process in triggering and perpetuating AF ^16^. Inflammatory mediators cause increased damage in the atrial myocardium, altering electrical pathways and driving atrial fibrosis, thereby forming major substrates that promote AF persistence ^17^. Clinical findings have revealed that circulating proteins such as CRP, interleukin-6 and tumour necrosis factor-alpha are higher in plasma from AF patients ^16^, with CRP levels particularly elevated in persistent AF (persAF) compared with paroxysmal AF (pxAF) patients ^18^. This is further supported by laboratory evidence showing that inflammatory responses are prominent not only at the systemic level, but also locally in the atria, targeting cardiomyocytes and cardiac fibroblasts ^19^. Circulating inflammatory markers such as matrix metalloproteinase 2, neurogenic locus notch homologue protein 3, and tumour necrosis factor-receptor 2 are involved in atrial inflammatory responses and associated with longer AF episode duration ^12^. Cardiac ablation is the standard surgical procedure for AF, which can suppress chronic inflammation and reverse pathological remodelling in the atria upon successful ablation ^20^. While it has been suggested that matrix metalloproteinases contribute to this remodelling process ^20^, further understanding on the molecular mechanisms involved beyond inflammatory pathways is required.

This study aims to identify new plasma protein signatures of AF in an exploratory cohort using the highly sensitive Olink assay. In contrast to previous studies using Olink Cardiovascular panels with only 92 proteins ^21,22^, we employed the Olink Reveal panel, which offers broader protein coverage of inflammatory and immune response processes with 1034 proteins. Profiling of plasma proteomes from patients at both baseline and post-ablation potentially reveals novel molecular targets contributing to AF-associated inflammatory responses in the underlying pathophysiology.

## Methods

### Study design and patient population

This study was conducted under the approval by the South Central-Berkshire B Research Ethics Committee (UK, 18/SC/0304) at the John Radcliffe Hospital, Oxford. Patients who were admitted for catheter ablation, cardioversion or routine electrophysiology study were recruited to this study and provided written informed consent before enrolment. Two disease groups were classified: 1) pxAF, defined as patients with AF episodes that stop within 7 days, and 2) persAF, patients with AF episodes longer than 7 days. The non-AF control group included patients with supraventricular tachycardias or atrial flutter, without AF history. Diagnosis of AF and non-AF arrhythmic events were identified using electronic patient records, ECG measurements and electrophysiology study. Thirty patients were recruited in total, including ten non-AF controls, ten pxAF patients and ten persAF patients. Demographic and clinical characteristics from patients were retrieved from the NHS electronic patient records by qualified clinicians and researchers in the study team. Biochemical measurements were recorded when available to assess changes in physiological functions. Additionally, to further assess cardiac structural and functional outcomes of the patients, electrocardiogram (ECG) measurements and echocardiography parameters were also collected when available.

### Blood collection and plasma processing

Blood samples were collected from patients who visited the hospital for the procedure (baseline – obtained before ablation) and returned for a follow up visit 12 months after the procedure. Specifically, peripheral venous blood samples were collected from the femoral veins at baseline, and from the antecubital veins at the follow up visit 12 months after the procedure. Blood was collected using sterile syringes by a member of the study’s clinical team. After collection, the samples were immediately transferred to BD Vacutainer Citrate Tubes and left to clot at room temperature for 1 hour. The tubes were then centrifuged at 3000 rpm for 15 minutes at room temperature for layer separation. Plasma from the top layer was extracted and kept at −80°C for long-term storage.

### Processing of samples and data for Olink assay

Protein levels in human plasma were measured using the Olink Reveal panel (Olink Proteomics AB, Uppsala, Sweden) using the proximity extension assay technology. 40 µl of plasma was loaded to each well on the clean 96-well PCR-plate with full skirt (#AB0800, ThermoFisher Scientific). Plate was sealed with MicroAmp seal (#4306311, ThermoFisher Scientific), stored at −80°C and kept on dry ice during transfer to the Multiomics Technology Platform Group (Oxford, UK) for the generation and initial processing of the sequencing data. Samples were sequenced by next generation sequencing using Illumina’s NovaSeq 6000 platform. Protein levels were quantified and expressed as the Olink’s Normalised protein expression (NPX) values.

### Statistical analysis

Statistical analysis was conducted using GraphPad Prism 10.5.0 and R version 4.3.2. Normal distribution was determined by Shapiro-Wilk test. For multi-group comparisons, normally distributed data were analysed by one-way analysis of variance (ANOVA); non-parametric data – by Kruskal-Wallis test. Differences between multiple categorical variables were analysed using Fisher’s exact test. Differential expression analysis was performed using the package *limma* (v3.58.1) in R. NPX values from Olink were used for analysis, followed by fitting linear models to each protein using *limma*, and applying empirical Bayes moderation to stabilise variance estimates. For the baseline analysis, differences were assessed using a model including the disease group conditions only. For analyses of 12-month follow-up samples, a paired design was implemented by including donor (patient) as an adjusting factor in the model, and hence, accounting for within-individual differences. Moderated t-statistics and p-values were reported in the results. Given the exploratory nature of this study, differentially expressed proteins (DEPs) were determined using a non-adjusted *P*-value < 0.05.

Binary outcomes were modelled using Firth’s bias-reduced logistic regression and this was analysed using the package *logistf* in R (v1.26.1). Given the small sample size and low number of events, Firth’s penalised likelihood approach was implemented over the conventional logistic regression. Continuous outcomes were analysed using Ridge linear regression with the package *glmnet* (v.4.1-9) in R. Ridge regression was applied instead of ordinary least-squares linear regression due to limited sample size. Data visualisation and graphs (volcano plots, forest plots, heatmaps, dot plots, bar plots) were generated using the package *ggplot2* (v3.5.2) in R. *P*-values less than 0.05 were statistically significant.

## Results

### Baseline patient characteristics

Proteome profiling was carried out on plasma blood samples collected from a total of thirty patients (**Fig. 1a**) who underwent catheter ablation. Analysis of the baseline demographic and clinical characteristics (**Table 1**) shows that the use of anticoagulants was more prevalent in patients with pxAF (90%) and persAF (100%) compared to non-AF controls (50%) (*P* = 0.0265). Baseline levels of urea were higher in persAF (6.7 ± 1.4 mmol/L) than pxAF (5.1 ± 1.3 mmol/L) and non-AF (5.3 ± 1.4 mmol/L) (*P* = 0.0234); however, the prevalence of reduced estimated glomerular filtration rate (eGFR) was similar between the groups (*P* = 0.9999). Baseline ECG measurements indicated a higher heart rate in persAF patients (87.0 ± 21.1 beats per minute [bpm]) compared with the other two groups (non-AF: 65.4 ± 12.8 bpm; pxAF: 70.4 ± 11.5 bpm). Other characteristics did not differ between groups.

**Figure 1.**
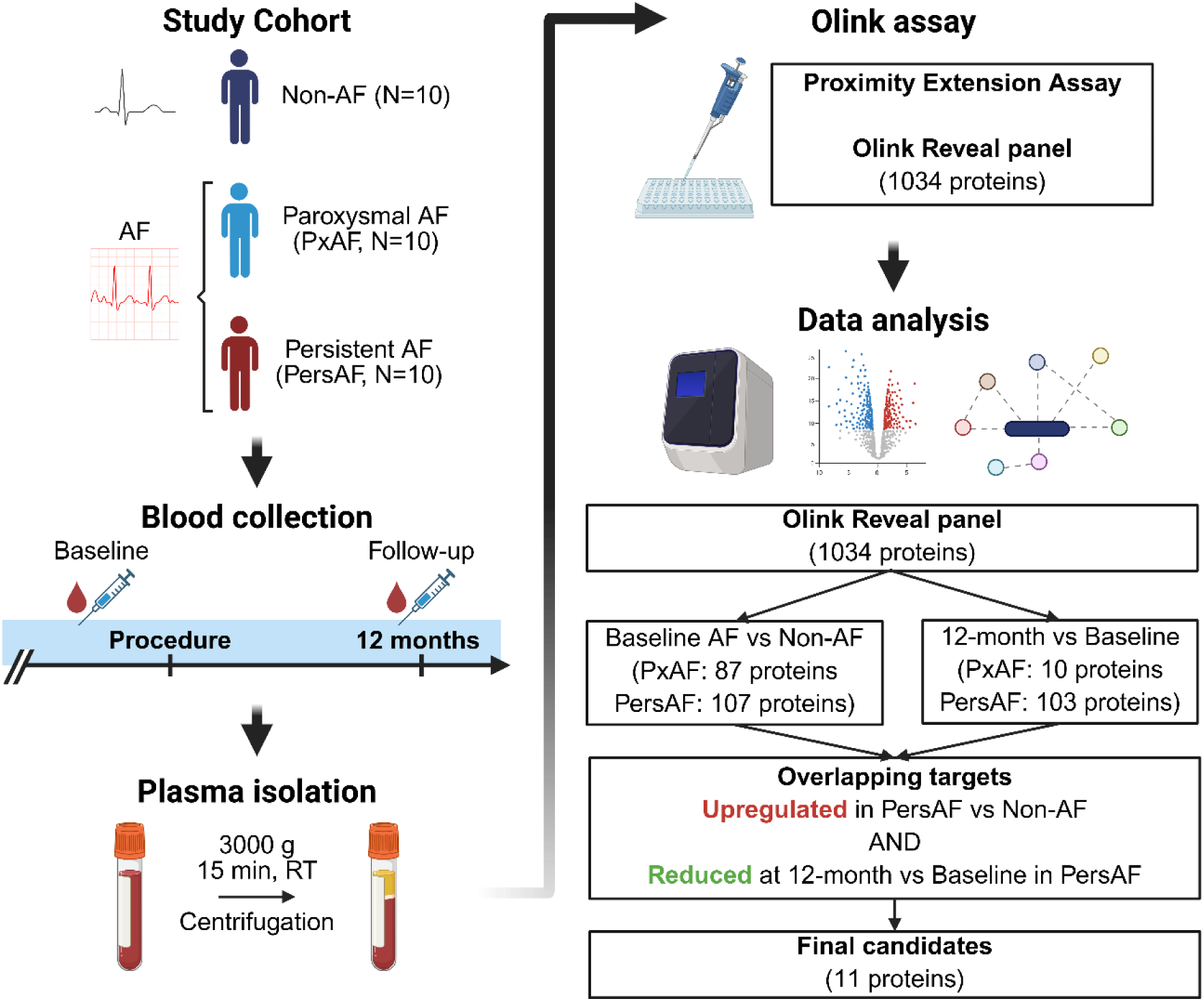
Study design and data analysis workflow. **(left)** Experimental design showing study cohort, blood sample collection and plasma isolation. **(right)** Olink assay and data analysis workflow, identifying differentially expressed proteins and shortlisting the 11 candidate proteins associated with persistent AF. Created with BioRender.com.

**Table 1.** Baseline demographic and clinical characteristics of patients.

| Characteristic | Non-AF<br>(n=10) | PxAF<br>(n=10) | PersAF<br>(n=10) | P-value |
| --- | --- | --- | --- | --- |
| **Demographic** |  |  |  |  |
| Male, n (%) | 6 (60) | 7 (70) | 8 (80) | 0.8792 |
| Age, y (SD) | 58.0 ± 12.7 | 62.5 ± 9.1 | 61.1 ± 7.1 | 0.5882 |
| Body mass index, kg/m<sup>2</sup> (SD) | 29.2 ± 6.2 | 29.0 ± 4.6 | 28.9 ± 5.9 | 0.9546 |
| Current/ex-smoker, n (%) | 1 (10) | 2 (20) | 5 (50) | 0.1916 |
| Alcohol consumption, n (%) | 6 (60) | 9 (90) | 4 (40) | 0.0891 |
| **Ablation procedure, n (%)** |  |  |  |  |
| Radiofrequency ablation | 9 (90) | 5 (10) | 9 (90) | 0.1512 |
| Cryoablation | 1 (10) | 4 (40) | 1 (1) | 0.3001 |
| Cardioversion | 0 (0) | 3 (30) | 5 (50) | 0.0527 |
| **Medical history, n (%)** |  |  |  |  |
| Ischemic heart disease | 1 (10) | 1 (10) | 1 (10) | 0.9999 |
| Stroke/TIA | 0 (0) | 0 (0) | 1 (10) | 0.9999 |
| Myocardial infarction | 2 (20) | 0 (0) | 0 (0) | 0.3103 |
| Hypertension | 1 (10) | 3 (30) | 3 (30) | 0.6419 |
| Peripheral arterial disease | 0 (0) | 0 (0) | 0 (0) | 0.9999 |
| Diabetes | 0 (0) | 0 (0) | 0 (0) | 0.9999 |
| COPD | 0 (0) | 0 (0) | 0 (0) | 0.9999 |
| Chronic kidney disease | 0 (0) | 0 (0) | 0 (0) | 0.9999 |
| Heart failure | 2 (20) | 0 (0) | 1 (10) | 0.7537 |
| Thyroid disease | 0 (0) | 0 (0) | 0 (0) | 0.9999 |
| **Medication, n (%)** |  |  |  |  |
| Anticoagulant | 5 (50) | 9 (90) | 10 (100) | 0.0265 |
| Antiplatelet | 1 (10) | 1 (10) | 0 (0) | 0.9999 |
| β-blocker | 6 (60) | 6 (60) | 7 (70) | 0.9999 |
| Amiodarone | 0 (0) | 1 (10) | 2 (20) | 0.7537 |
| Flecainide | 2 (20) | 3 (30) | 0 (0) | 0.3211 |
| Statins | 5 (50) | 4 (40) | 2 (20) | 0.5101 |
| Calcium channel blocker | 1 (10) | 3 (30) | 2 (20) | 0.8465 |
| Digoxin | 0 (0) | 0 (0) | 2 (20) | 0.3103 |
| Nitrates | 0 (0) | 1 (10) | 0 (0) | 0.9999 |
| ACEi | 3 (30) | 4 (40) | 3 (30) | 0.9999 |
| Diuretics | 2 (20) | 2 (20) | 2 (20) | 0.9999 |
| **Blood measurements** |  |  |  |  |
| Haemoglobin, g/L (SD) | 136.9 ± 21.8 | 149.3 ± 8.8 | 151.3 ± 13.3 | 0.1005 |
| White blood count, x 10<sup>9</sup>/L (SD) | 6.2 ± 1.3 | 7.1 ± 1.5 | 6.6 ± 1.5 | 0.3311 |
| Urea, mmol/L (SD) | 5.3 ± 1.4 | 5.1 ± 1.3 | 6.7 ± 1.4 | 0.0234 |
| Creatinine, μmol/L (SD) | 76.5 ± 18.8 | 75.4 ± 16.8 | 83.0 ± 17.2 | 0.5867 |
| Reduced eGFR<br>($\leq 89$ ml/min/1.73 m<sup>2</sup>), n (%) | 6 (60) | 5 (50) | 5 (50) | 0.9999 |
| **Cardiac parameters** |  |  |  |  |
| Reduced LVEF, n (%) | 4 (40) | 2 (20) | 5 (50) | 0.5101 |
| Heart rate, bpm (SD) | 65.4 ± 12.8 | 70.4 ± 11.5 | 87.0 ± 21.1 | 0.0260 |
| QT interval, ms (SD) | 415.5 ± 31.2 | 435.1 ± 34.3 | 430.7 ± 33.7 | 0.2042 |
| QRS duration, ms (SD) | 92.6 ± 19.3 | 90.1 ± 7.9 | 97.4 ± 15.1 | 0.5434 |

ACEi, angiotensin-converting enzyme inhibitor; bpm, beats per minute; COPD, chronic obstructive pulmonary disease; eGFR, estimated glomerular filtration rate; LVEF, left ventricular ejection fraction; PersAF, persistent atrial fibrillation; PxAF, paroxysmal atrial fibrillation; TIA, transient ischemic attack. *P*-values calculated by one-way ANOVA (Age, haemoglobin, white blood count, urea, creatinine, QRS duration), Kruskal-Wallis test (body mass index, heart rate, QT interval), and Fisher's exact test for other characteristics.

### Differentially expressed proteins in pxAF and persAF at baseline

Proteins in the Olink Reveal panel were analysed in the downstream differential expression analysis. We identified DEPs specific to pxAF and persAF compared with non-AF control patients. In the pxAF group, 82 upregulated and 5 downregulated proteins were identified (**Fig. 2a** and **Supplementary Table S1**). In the persAF group, 101 upregulated and 6 downregulated proteins were identified (**Fig. 2b** and **Supplementary Table S2**). The top 5 upregulated and top 5 downregulated DEPs are visualised in the forest plots for pxAF (**Fig. 2c**) and persAF (**Fig. 2d**).

**Figure 2.**
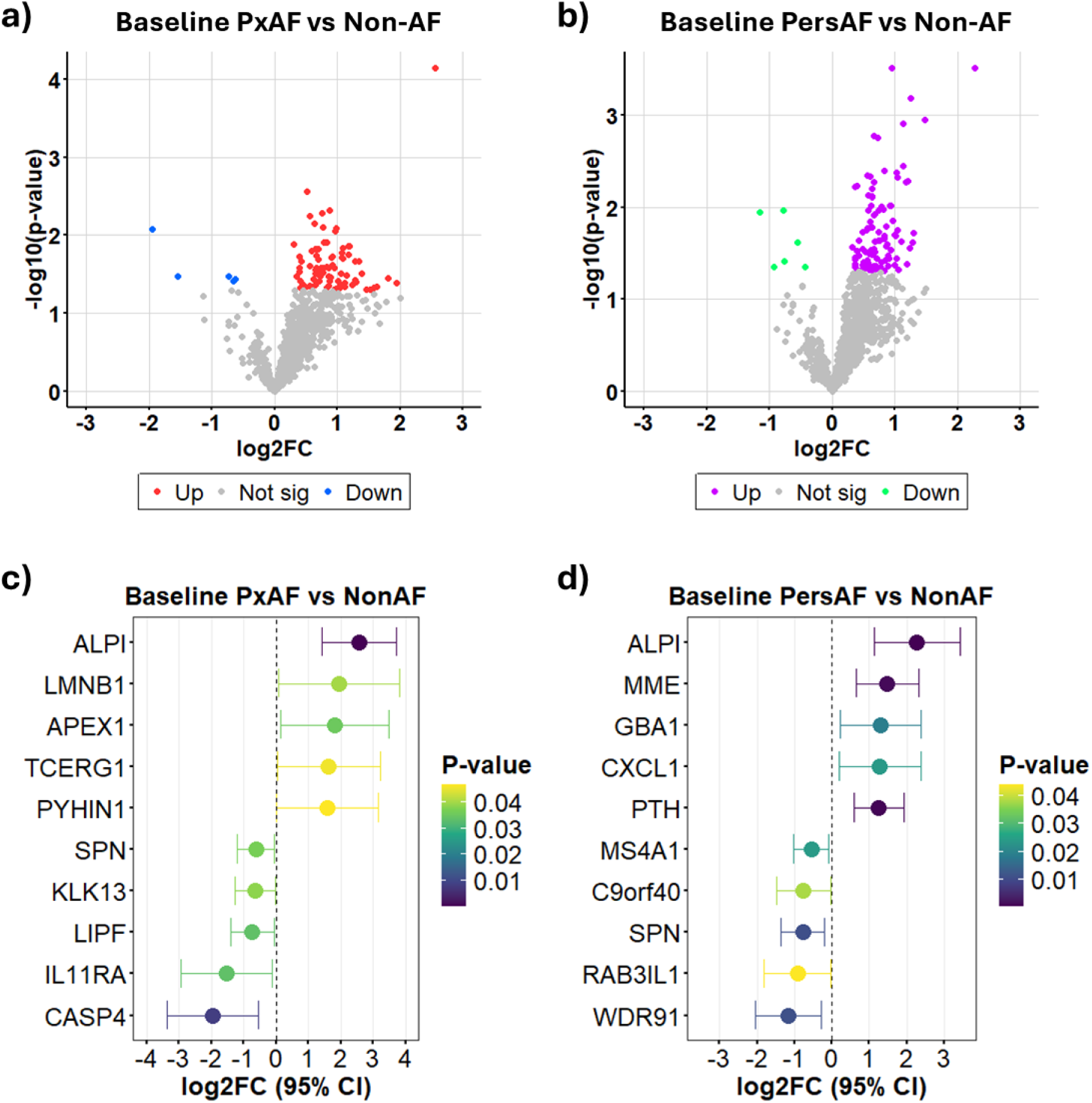
Differentially expressed proteins in paroxysmal and persistent atrial fibrillation. **(a)** Volcano plot showing proteins compared between pxAF and non-AF patients. Significant proteins (*P* < 0.05) are highlighted: upregulated in red and downregulated in blue. **(b)** Volcano plot showing proteins compared between persAF and non-AF patients. Significant proteins (*P* < 0.05) are highlighted: upregulated in purple and downregulated in light green. **(c)** Forest plot showing top 5 positively and top 5 negatively regulated proteins in pxAF patients. **(d)** Forest plot showing top 5 positively and top 5 negatively regulated proteins in persAF patients. Plasma protein levels were expressed as NPX values from the Olink assay. Differential analysis was performed using *limma*.

### Differentially expressed proteins in AF patients at 12 months post-ablation

In each study participant with AF, plasma proteome was profiled at baseline (before ablation) and 12 months post-ablation. In AF patients (both pxAF and persAF), 9/20 (45%) developed AF recurrence within 12 months after ablation. Differential analysis revealed 5 upregulated and 5 downregulated DEPs at 12 months after ablation in pxAF patients (**Fig. 3a** and **Supplementary Table S3**). In contrast to pxAF, more DEPs were identified in patients with persAF post-ablation, including 69 upregulated and 34 downregulated proteins (**Fig. 3b** and **Supplementary Table S4**). The top 5 upregulated and top 5 downregulated DEPs are visualised in the forest plots for post-ablation changes in pxAF (**Fig. 3c**) and persAF (**Fig. 3d**). The plasma proteome in non-AF patients post-ablation was also profiled (**Supplementary Table S5**).

**Figure 3.**
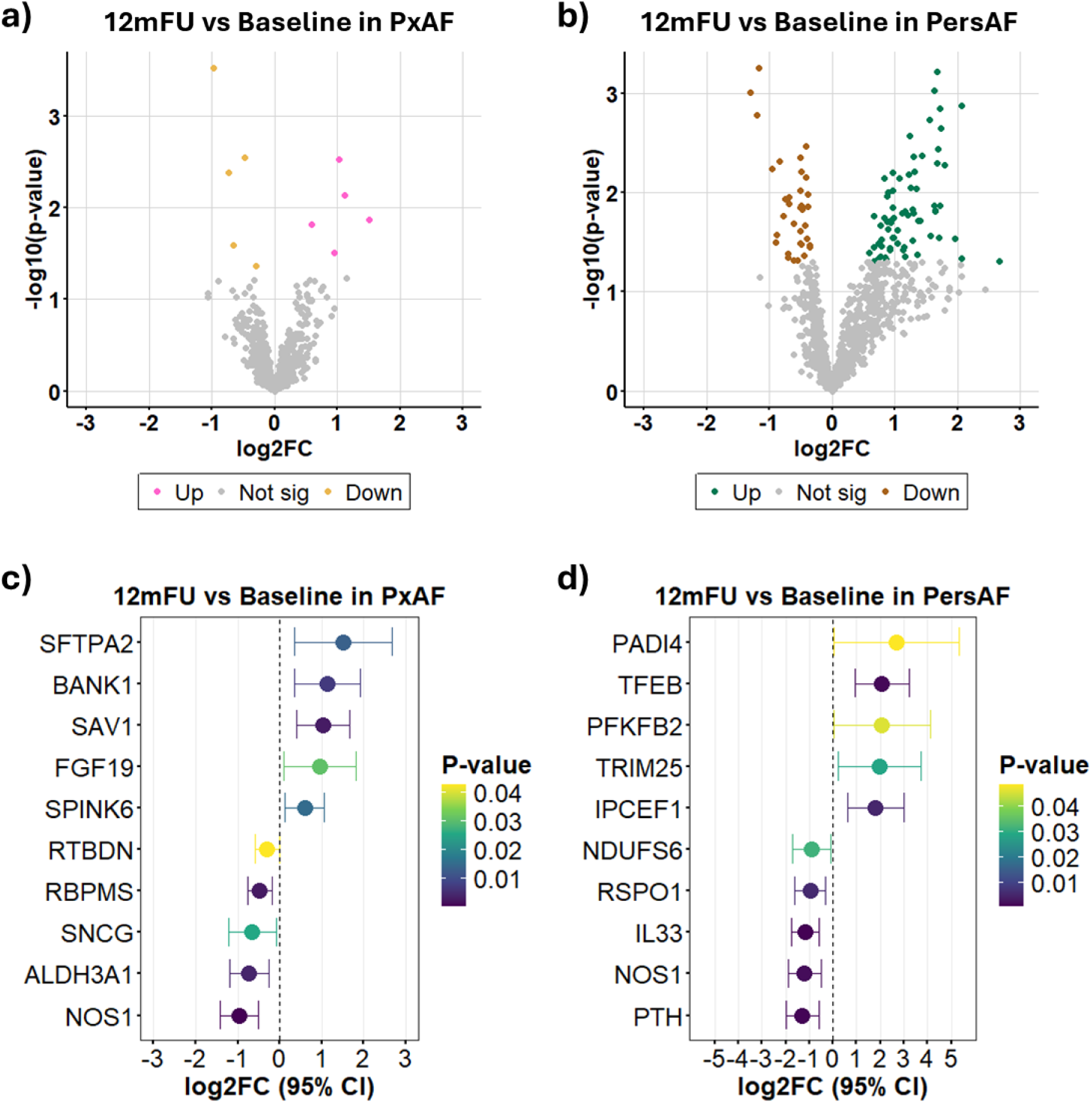
Differentially expressed proteins in atrial fibrillation patients at 12 months post-ablation. **(a)** Volcano plot showing protein levels in pxAF patients between baseline and 12 months follow-up (12mFU). Significant proteins (*P* < 0.05) are highlighted: upregulated in pink and downregulated in yellow. **(b)** Volcano plot showing protein levels in persAF patients between baseline and 12mFU. Significant proteins (*P* < 0.05) are highlighted: upregulated in dark green and downregulated in brown. **(c)** Forest plot showing top 5 positively and top 5 negatively regulated proteins in pxAF patients at follow-up. **(d)** Forest plot showing top 5 positively and top 5 negatively regulated proteins in persAF patients at follow-up. Plasma protein levels were expressed as NPX values from the Olink assay. Differential analysis was performed using *limma*.

### Selection of candidate proteins associated with pxAF and persAF

To shortlist candidate proteins involved in AF pathophysiology in relation to the ablation procedure, we applied a selection strategy based on protein expression patterns associated with both AF status (pxAF and persAF) at baseline and its changes over time – post-ablation. We selected candidate targets based on either of two criteria: (1) upregulated DEPs at baseline that were decreased 12 months after ablation, or (2) downregulated DEPs at baseline that were increased post-ablation. This strategy may reflect underlying biological processes responsive to rhythm restoration by ablation.

In pxAF patients, we first identified 87 proteins that were differentially expressed between the pxAF and non-AF groups at baseline. Among these, only 1 protein (ALDH3A1) demonstrated a significant directional shift 12 months after ablation, where it was elevated in pxAF at baseline and showed reduced expression post-ablation (**Fig. 4a-4b**). The strategy was also used in the persAF group, which identified 11 DEPs at baseline that exhibited a significant directional shift 12 months after ablation. Specifically, 11 proteins that were upregulated in persAF at baseline showed reduced expression 12 months after ablation (**Fig. 4c**), whereas no proteins were recognised with the opposite pattern (**Fig. 4d**). Accordingly, these 11 proteins were identified as candidates demonstrating partial normalisation of expression following rhythm restoration (**Fig. 4e**). Baseline expression levels of these candidates across groups are visualised in a heatmap (**Fig. 4f**), and their corresponding post-ablation changes in persAF patients are subsequently illustrated (**Fig. 4g**).

**Figure 4.**
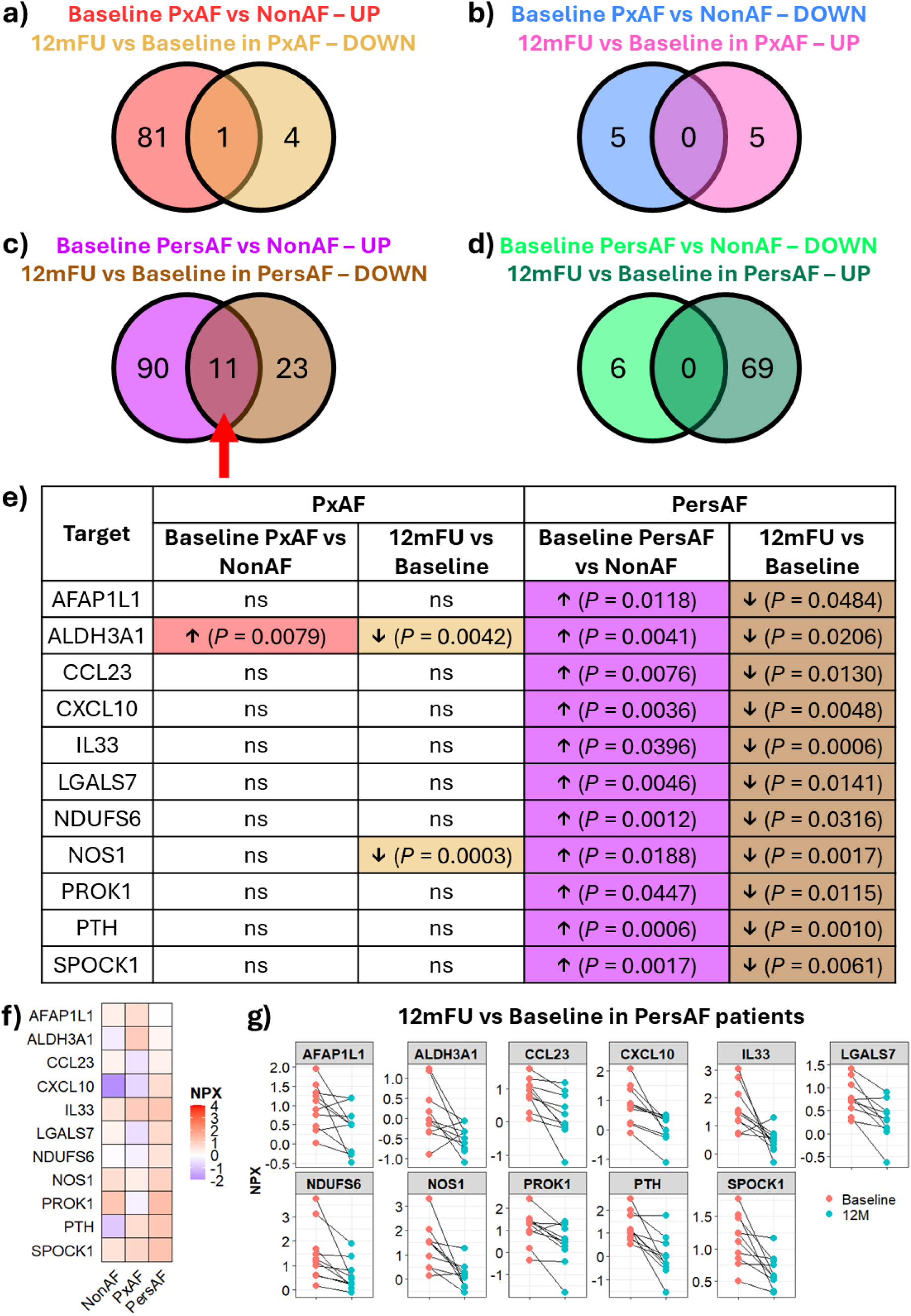
Selection of 11 candidate proteins associated with atrial fibrillation. Differential analyses for disease vs control and follow-up vs baseline were integrated. **(a-d)** Venn diagrams showing number of proteins from the corresponding differential analysis and number of overlapping proteins (intersection) between the two comparisons. Red arrow pointing the 11 proteins to be shortlisted as candidate proteins associated with AF. **(e)** Table summary of the regulation direction and *P*-values of the candidate proteins in pxAF and persAF patients. **(f)** Heatmap showing baseline NPX levels of the candidate proteins. **(g)** Dot plots showing the changes of the candidate protein levels between baseline and 12 months follow-up after ablation (12mFU). Line and paired dots indicating changes in the same donor.

### Association between AF-related proteins and baseline clinical features

We assessed the interrelationships of the 11 candidate proteins using correlation analysis, and all significant protein pairs showing positive correlations (correlation coefficient > 0, *P* < 0.05; **Fig. 5a** and **Supplementary Table S6**). Proteins such as AFAP1L1, CXCL10, NOS1 and SPOCK1 showed moderate-to-strong positive correlations with most other candidates, indicating potential co-regulation or shared upstream biological pathways.

**Figure 5.**
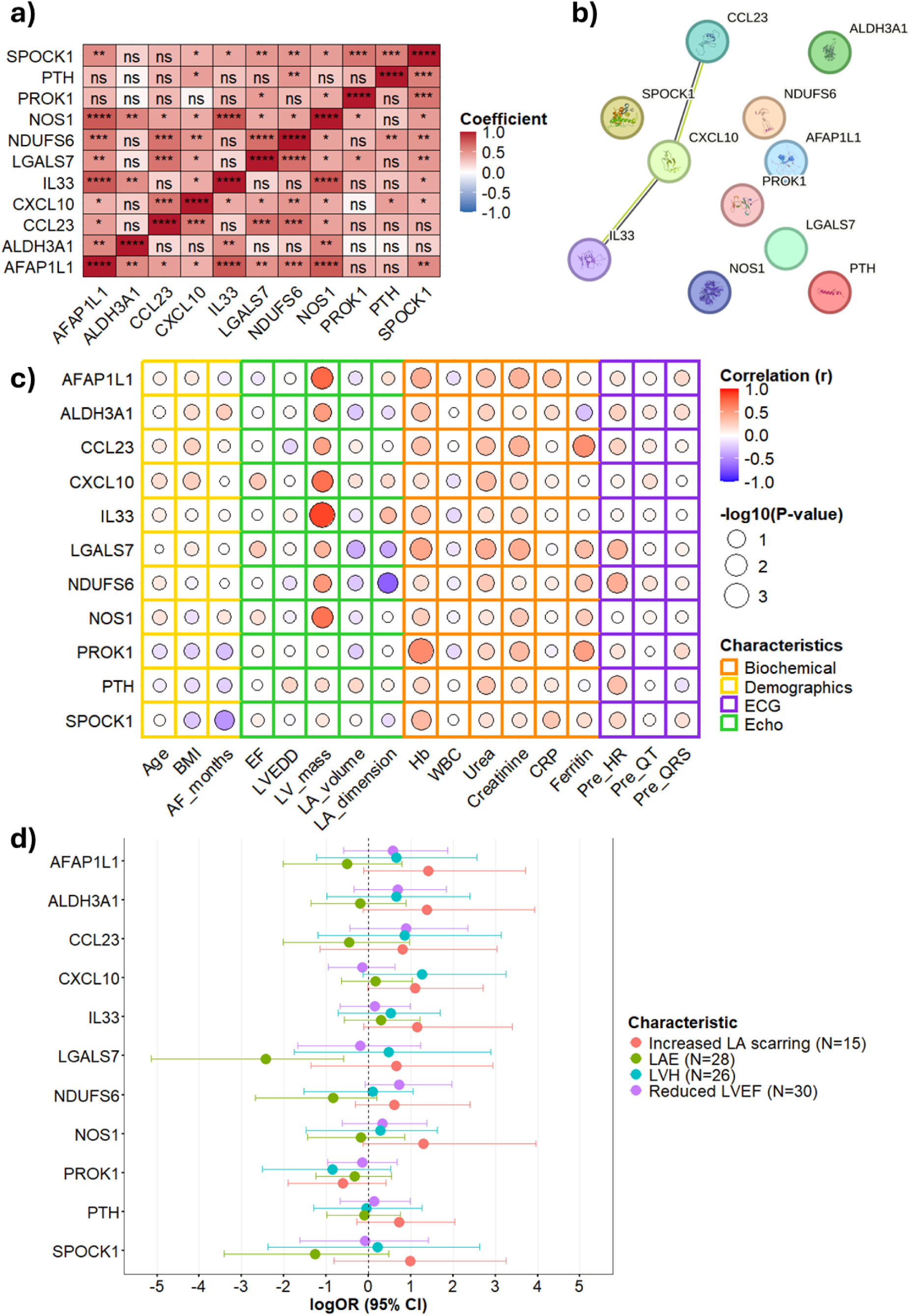
Associations of candidate proteins with baseline clinical features. **(a)** Heatmap showing a correlation matrix of the baseline expression levels between the candidate proteins. \**P* < 0.05, \*\**P* < 0.01, \*\*\**P* < 0.001, \*\*\*\**P* < 0.0001, ns = not significant. **(b)** Protein-protein interaction (PPI) network of the candidate targets generated using the STRING database (medium confidence 0.400). **(c)** Heatmap showing a correlation matrix between the protein expression and baseline clinical features. Features are categorised and highlighted in different colours: biochemical (red), demographics (yellow), ECG (purple) and echocardiography (green). **(d)** Forest plot showing the associations of baseline protein levels with characteristics related to cardiac remodelling. Log odds ratio (logOR) was calculated using univariate Firth’s logistic regression.

In contrast, protein pairs involving ALDH3A1, PROK1, or PTH largely demonstrated non-significant correlations, suggesting that a subset of these proteins may function independently in AF. To obtain a broader view of the interactions among these proteins, a protein-protein interaction network was generated using the STRING database (**Fig. 5b**). Evidence of interaction was identified in two protein pairs: CCL23-CXCL10 and CXCL10-IL33. Together, the correlation analysis and protein-protein interaction network suggest that these candidate proteins do not act as a single molecular player but instead represent a combination of interconnected and independent biological signals that may contribute to AF pathogenesis, particularly in patients with persAF.

We next explored the clinical relevance of the 11 candidate proteins by assessing their correlations with demographic variables, biochemical markers, ECG parameters and echocardiographic measurements (**Fig. 5c** and **Supplementary Table S7**). The results revealed a heterogenous patterns of associations between protein expression levels and clinical characteristics. Several proteins showed strong positive correlations with indexed left ventricular (LV) mass, a key indicator of LV hypertrophy (LVH) ^23^. A subset of candidates also displayed moderate positive correlations with circulating levels of haemoglobin and creatinine. Proteins associated with structural or functional echocardiographic parameters may reflect pathways associated with cardiac remodelling in AF, whereas correlations with biochemical markers may indicate systemic or metabolic influences in AF. In contrast, proteins with minimal associations may reflect biological processes not directly captured by the currently available clinical measurements or may reflect limited statistical power due to the exploratory cohort size, warranting further validation in larger studies.

To better understand the associations of these proteins with cardiac remodelling at baseline, we performed Firth’s bias-reduced logistic regression analyses examining the relationship between each protein and different remodelling phenotypes (**Fig. 5d** and **Supplementary Table S8**): increased left atrial (LA) scarring indicated by higher Utah stages (stage ≥ 2), LA enlargement (LAE), LVH and reduced LV ejection fraction (LVEF). Although most targets demonstrated no statistically significant associations with the phenotype, several showed noteworthy trends. For atrial remodelling, higher baseline LGALS7 levels were associated with lower odds of LAE (logOR −2.4308 [−5.1389, −0.5952], *P* = 0.0067). Several proteins also showed positive trends towards greater severity in LA scarring, including CXCL10 (logOR 1.1037 [−0.0102, 2.7036], *P* = 0.0523), AFAP1L1 (logOR 1.4062 [−0.1072, 3.7073], *P* = 0.0710), ALDH3A1 (logOR 1.3835 [−0.1321, 3.9221], *P* = 0.0765), IL33 (logOR 1.1557 [−0.1138, 3.3938], *P* = 0.0782) and NOS1 (logOR 1.3042 [−0.1234, 3.9562], *P* = 0.0797). In the context of ventricular remodelling, CXCL10 showed a positive association with LVH (logOR 1.2713 [−0.1278, 3.2472], *P* = 0.0782) while higher NDUFS6 levels were associated with increased odds of reduced LVEF (logOR 0.6055 [−0.0865, 1.9665], *P* = 0.0823).

### Predictive value of pxAF- and persAF-associated proteins for post-ablation and long-term outcomes

The electrophysiological relevance of the baseline levels of the shortlisted proteins for both post-ablation and long-term outcomes was assessed using ridge linear regression models. The contributions of each protein to post-ablation and 12 months follow-up ECG parameters (heart rate, QT interval and QRS duration) were presented as standardised coefficients across the outcomes. For heart rate, baseline CCL23 emerged as the top positive contributor in the post-ablation analysis (**Fig. 6a**), and consistently at 12 months follow-up as the second strongest positive contributor (**Fig. 6b**). This suggests that higher baseline levels of CCL23 relative to other proteins are more associated with higher heart rates post-ablation and at long-term follow-up. For the QT interval, baseline LGALS7 and SPOCK1 emerged as the two proteins with the most notable negative associations (**Fig. 6c-6d**), as their negative coefficients became more pronounced at 12 months follow-up compared to the other candidate proteins. This indicates a possible inverse relationship between baseline LGALS7 and/or SPOCK1 levels and QT duration in the long-term post-ablation period. For QRS duration, no coherent directional pattern was observed between post-ablation and 12 months follow-up analyses (**Fig. 6e-6f**).

**Figure 6.**
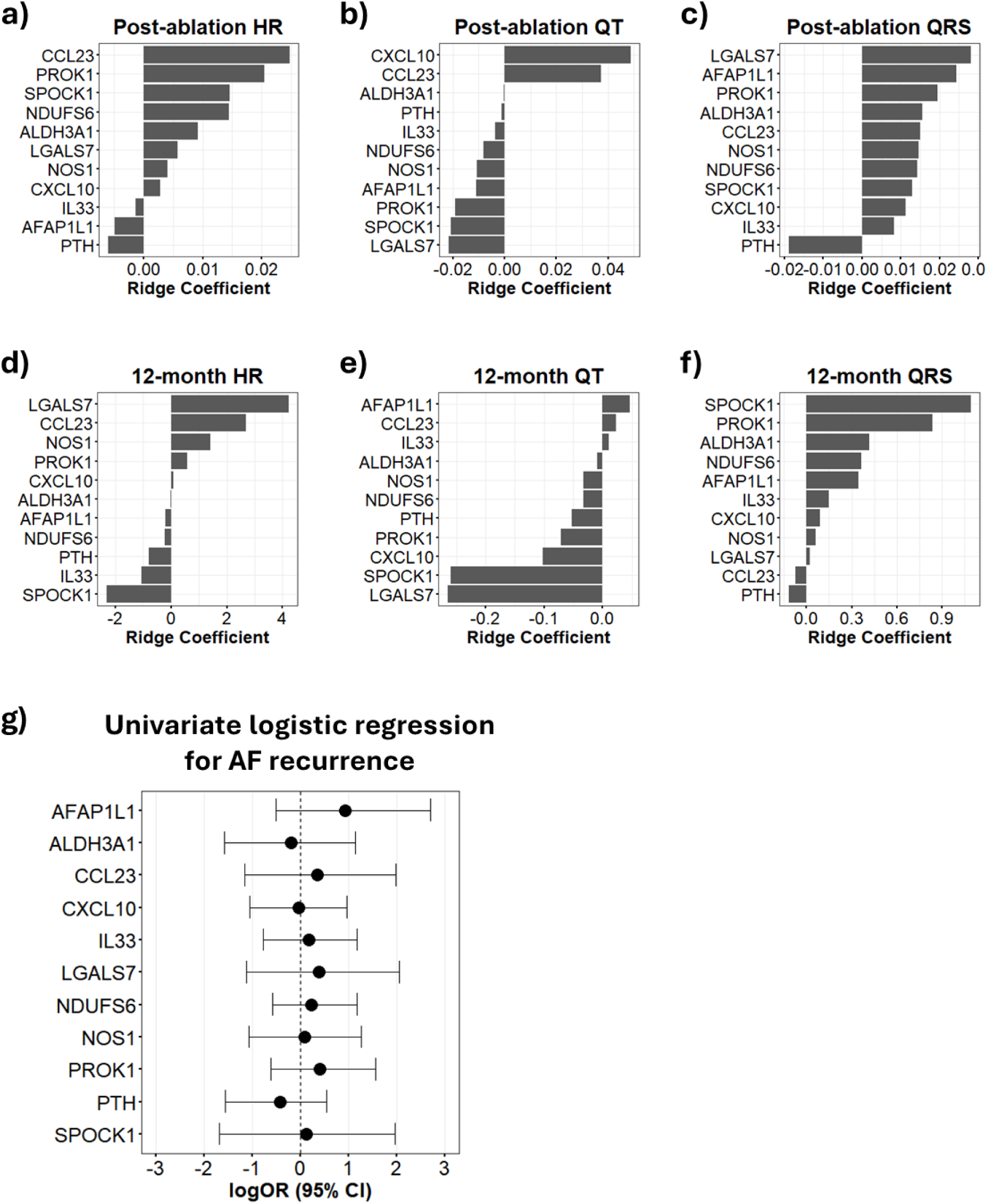
Associations of baseline levels of candidate proteins with immediate and 12 months post-ablation outcomes. Contributions of baseline levels of each protein to ECG outcomes were analysed using Ridge’s regression. **(a-c)** Bar plots showing standardised coefficients relating to post-ablation heart rate, QT interval and QRS duration. **(d-f)** Bar plots showing standardised coefficients relating to 12 months follow-up heart rate, QT interval and QRS duration. **(g)** Forest plot showing the associations of baseline protein levels with AF recurrence within 12 months post-ablation. Log odds ratio (logOR) was calculated using univariate Firth’s logistic regression.

To evaluate whether baseline levels of the 11 candidate targets were associated with AF recurrence within 12 months post-ablation, we performed Firth’s logistic regression for all AF patients (pxAF and persAF), fitting each protein as a univariate model (**Fig. 6g** and **Supplementary Table S9**). None of the proteins showed statistically significant association with AF recurrence. These findings suggest that while the shortlisted proteins were linked to AF status, their baseline levels alone do not predict AF recurrence over a 12-month follow-up.

## Discussion

In this study, we established a small exploratory cohort of patients who underwent catheter ablation and profiled plasma protein expression at baseline and 12 months post-ablation. We demonstrated distinct protein signatures in pxAF and persAF patients and evaluated the long-term regulation of these proteins after ablation.

The clinical hallmark of AF, characterised by higher heart rates, was reflected in the ECG measurements of our cohort, demonstrating a representative phenotype of AF in our disease groups. There was significantly higher use of anticoagulants in patients with pxAF and persAF than in non-AF patients at baseline. This was expected, as anticoagulant agents are crucial to reduce the risk of stroke in AF ^24^. We also identified higher urea levels specifically in the persAF group compared with non-AF and pxAF individuals. However, renal function was comparable among all three groups, as indicated by similar levels of creatinine and eGFR.

Our differential analysis of plasma proteins showed significantly more upregulated than downregulated proteins in pxAF and persAF at baseline. This could be explained by the composition of the Olink Reveal panel, which includes a majority of inflammatory protein markers. Given that systemic inflammation is a clear feature of AF, including in patients who have undergone catheter ablation ^25^, this pattern may reflect the predominance of upregulated over downregulated proteins in our plasma samples. Of particular interest, we observed differences in differentially regulated proteins between pxAF and persAF patients comparing with non-AF patients. This was not surprising, as the underlying proarrhythmic substrates and remodelling processes are markedly different between these phenotypes ^26^. PxAF involves earlier stages of AF progression predominantly characterised by electrical remodelling, whereas persAF represents later and more advanced stages with structural remodelling and an accumulation of fibrosis ^26^.

The ALPI protein was identified as the top upregulated protein in both pxAF and persAF patients. Elevation of circulating ALPI above physiological levels is commonly observed in liver-related conditions, such as cholestasis, cirrhosis and hepatitis ^27,28^. Alkaline phosphatase (AP) is closely linked to adverse outcomes in cardiovascular diseases; for example, high levels of AP have been associated with an elevated risk of mortality in coronary heart disease ^29^ and with cardiovascular events in chronic kidney disease ^30^. These associations have also been seen in patients with AF, where high serum AP levels were significantly associated with an elevated risk of cardiovascular events and hospital admission for heart failure ^31^. Although a study reported no significant difference in circulating AP levels between patients with and without AF at baseline, a trend was observed (*P* ≥ 0.083), and that cohort was restricted to male patients only ^32^. These findings highlight the prognostic and potential therapeutic value of ALPI in cardiovascular diseases and measuring its circulating level may help identify high-risk patients who may develop AF.

The 11 selected AF-associated candidate proteins underline the multifactorial pathogenesis of AF, reflecting a combination of four classes of biological functions. First, mediators of immune response and inflammation (CCL23, CXCL10, IL33) point to an active pro-inflammatory state, which is a major driver of electrical dysregulation and atrial fibrosis in AF ^16,17^. This immunomodulation and inflammatory signalling may facilitate structural reorganisation of the extracellular matrix and cell cytoskeleton ^33^, which includes the second class of proteins with cell adhesion and structural organisation properties (AFAP1L1, LGALS7, and SPOCK1). The excessive accumulation and maturation of extracellular matrix components in the tissue microenvironment form physical substrates that promote atrial fibrosis and sustain AF ^34^. The third class of proteins is involved in metabolic pathways essential for balancing oxidative stress (ALDH3A1) and cellular energy production in mitochondria (NDUFS6). Metabolic changes and impairment are key mechanisms promoting AF by disrupting cardiomyocyte contractions and driving oxidative stress-induced electrical and structural remodelling ^35^. The final class of proteins including NOS1, PROK1, and PTH represent regulation of physiological signalling and homeostasis. These proteins highlight critical processes that may alter electrical remodelling and atrial contraction, including PTH regulation of calcium homeostasis ^36^, NOS1 regulation of endogenous nitric oxide ^37^, and PROK1 regulation of a wide range of physiological functions such as angiogenesis, inflammation, and nociception ^38^. Collectively, these candidate proteins represent a holistic molecular profile integrating systemic inflammation and metabolic dysregulation that could exacerbate the fibrosis and electrophysiological dysfunction central to AF progression and maintenance.

This subset of candidate targets expressed strong positive correlations with LV mass (AFAP1L1, CXCL10, IL33 and NOS1). An increased LV mass is a clear clinical indication of LVH, and LVH is a known risk factor for AF incidence and is associated with a higher prevalence of comorbidities in AF, including diabetes, hypertension, and previous myocardial infarction ^39,40^. Although these studies presented the clinical association between the LVH phenotype and AF, our proteomic analyses in plasma from patients provide additional biological insights into specific molecular players that may be involved in this LVH-AF relationship. Also, among all baseline clinical features that were assessed, LV mass was exceptionally correlated with the baseline candidate proteins, which may imply that the biological effects of the plasma proteins are systemic or reflect global changes in the heart rather than being atrial-specific only. In addition to LV mass, moderate positive correlations between haemoglobin and the baseline candidates were observed (AFAP1L1, LGALS7, PROK1, SPOCK1). A retrospective cohort study demonstrated higher rates of AF incidence in patients with polycythaemia (high haemoglobin) ^41^. Polycythaemia contributes to pathophysiological outcomes by increasing hemodynamic shear stress, leading to conditions of pressure overload and adverse remodelling in the heart ^42,43^. Hence, these plasma proteins that correlated significantly with haemoglobin in AF patients may contribute to the underlying hemorheological risks as a mechanism to promote AF. However, these findings should be further studied and validated in larger cohorts.

While this study was directed to the plasma proteome in AF, regulation of the candidate proteins in the atria would be informative of their biological roles locally in the heart beyond systemic effects. Our previous analysis of a bulk RNA-sequencing public dataset from human left atrial appendages demonstrated transcriptional regulation of target genes in patients with persAF ^44,45^. Among the candidates, *CXCL10* gene expression was upregulated in persAF, consistent with the elevated CXCL10 protein levels in plasma from persAF patients. The concordant increase of CXCL10 in circulating and tissue levels may suggest a global AF-associated response rather than a systemic upregulation only. Moreover, while the plasma levels of NOS1 were elevated in persAF, gene expression of *NOS1* was significantly downregulated in the atria from persAF patients ^44,45^. Neuronal nitric oxide synthase protein, encoded by the *NOS1* gene, was also depleted in human and goat AF due to an atrial-specific upregulation of microRNA-31 ^46^. Hence, this divergence of protein regulation in plasma and tissue levels in AF suggests a compartment-specific regulation of NOS1 in AF and it may be modulated by distinct mechanisms in the systemic circulation compared to the tissue microenvironment.

We attempted to analyse whether the candidate proteins could be associated with AF recurrence and other post-ablation complications, but the event rates were very low and unable to yield significant findings due to being underpowered. Enhancing the understanding of proteins and pathways in patients with AF post-ablation has important clinical implications. It can improve more personalised management strategies for long-term outcomes, particularly AF recurrence after rhythm restoration. AF recurrence can be contributed by clinical comorbidities like hypertension, obesity and sleep apnoea, resulting in the need for repeat catheter ablation ^47^. Nevertheless, repeating ablation may induce extensive damage to the myocardium, exposing patients to a higher risk of surgical complications and other arrhythmic conditions, leading to abnormal contraction in the atria ^47,48^. Therefore, utilising protein markers may benefit in identifying patients at higher risk of recurrence and guide the optimal timing and selection of therapeutic interventions to prevent repeated ablation procedures.

Our study has several key strengths that enhance its scientific and clinical relevance. By profiling the plasma proteome at baseline and 12 months post-ablation, we uniquely capture dynamic molecular changes associated with rhythm restoration, a longitudinal design rarely explored in AF proteomics. The high-coverage Olink Reveal panel enabled broad interrogation of inflammatory, immune, metabolic, and signalling pathways involved in AF pathophysiology, providing deeper insight than smaller targeted panels. The identification of distinct proteomic signatures for pxAF and persAF underscores the biological heterogeneity of these phenotypes. This aligns with emerging evidence that circulating protein signatures can improve AF risk stratification and reflect underlying biological pathways such as extracellular matrix remodelling and inflammation ^15^. Correlation of candidate proteins with clinical features such as left ventricular mass and haemoglobin further integrates molecular and phenotypic dimensions of AF, and the partial normalisation of specific proteins after ablation suggests potential for future biomarker-guided stratification and monitoring.

Limitations of this study include that protein levels in the Olink assay were measured in NPX rather than exact concentrations. This restricts comparisons between different patients in different cohorts and makes it difficult to translate to clinical settings where absolute concentrations are necessary for making clinical decisions. Although this study was conducted with an exploratory nature, the sample size was limited and might be underpowered. Hence, adjusted *P*-values were not considered in differential analyses and covariates were not adjusted in our primary analyses to avoid the consequential loss of power. Also, we noticed a significantly higher use of anticoagulants in patients with AF, but this was not adjusted due to high collinearity between anticoagulants use and AF rhythm status. Finally, we acknowledge the existence of other large-scale studies presenting the plasma proteomes of AF patients. However, the results were limited to baseline samples and proteomic profiles from those patients after ablation were scarce. Our current investigation provides this additional layer of findings, informing the changes in protein regulation after undergoing ablation. Therefore, the identified AF proteomes and candidate proteins should be carefully considered as preliminary findings and require validation in external cohorts.

## Conclusion

In summary, this study identified plasma proteins differentially regulated in patients with AF. Notably, a subset of proteins was highlighted as candidates with increased expression in persAF at baseline and partially normalised after catheter ablation. These proteins imply important biological processes in AF pathophysiology and may serve as molecular markers for future studies investigating AF prediction models and therapeutic developments. It is critical to value the significance of plasma proteins in personalised diagnosis and treatment for patients with AF, in addition to the use of available electronic health records and existing biomarkers of AF.

## Supporting information

Supplementary Tables to be used for the link to the on the preprint site

## Abbreviations

AF: Atrial fibrillation
ANOVA: One-way analysis of variance
AP: Alkaline phosphatase
bpm: Beats per minute
CRP: C-reactive protein
DEP: Differentially expressed protein
ECG: Electrocardiogram
eGFR: Estimated glomerular filtration rate
LA: Left atrial
LAE: Left atrial enlargement
LV: Left ventricular
LVEF: Left ventricular ejection fraction
LVH: Left ventricular hypertrophy
NPX: Normalised protein expression
PersAF: Persistent atrial fibrillation
PxAF: Paroxysmal atrial fibrillation

## Acknowledgements

We are grateful for all patients who provided blood samples for this study, and all staff and investigators who contributed to this study. We thank the Multiomics Technology Platforms Group at the Centre for Human Genetics for the generation and initial processing of the sequencing data.

## Author contributions

C.H.K.Y., C.S.M. and S.R. conceived and designed the study. The clinicians T.R.B., K.R., M.G., M.P., Y.B. and R.W. collected and provided the blood samples from patients under the ethical approval granted to S.R.. C.H.K.Y. and C.S.M. performed the selection of samples and sample processing for Olink analysis. C.H.K.Y. contributed to the data processing and statistical analysis with the assistance of C.S.M.. The initial version of the manuscript was drafted by C.H.K.Y.; and C.H.K.Y., C.S.M. and S.R. jointly revised the manuscript. S.R. secured funding and supervised the work.

## Data availability statement

All data related to the analysis is included in the manuscript. The metadata of the patient cohort and raw values from Olink that support the findings are available from the first author (C.H.K.Y.) and corresponding author (S.R.) upon request.

